# Is it time to change the reference genome?

**DOI:** 10.1101/533166

**Authors:** Sara Ballouz, Alexander Dobin, Jesse Gillis

**Affiliations:** The Stanley Institute for Cognitive Genomics, Cold Spring Harbor Laboratory, Cold Spring Harbor, NY, 11724, USA

**Keywords:** Reference genome, consensus genomes, genetic diversity

## Abstract

The use of the human reference genome has shaped methods and data across modern genomics. This has offered many benefits while creating a few constraints. In the following piece, we outline the history, properties, and pitfalls of the current human reference genome. In a few illustrative analyses, we focus on its use for variant-calling, highlighting its nearness to a “type specimen”. We suggest that switching to a consensus reference offers important advantages over the current reference with few disadvantages.

## Why do we need references?

A block of platinum-iridium alloy sits in the International Bureau of Weights and Measures in France. Until May 20^th^, 2019, it will have a mass of precisely 1 kg. From then on, the kilogram (Le Grand K) will be defined in reference to Planck’s constant (6.626070150 x 10^−34^ kg·m^2^/s [1]) and this will not change for the foreseeable future. The human genomic location of the tumor protein p53 is chromosome 17: 7,666,487 −7,689,465 (genome reference GRCh38.p12). How permanent is the reference that determines this? We will never define the genome in terms of universal constants but can we do better than our current idiosyncratic choice?

### Frame of reference

We need standards to communicate in a common frame of reference, but not all standards are created equal. When the block of platinum-iridium loses a few atoms, the mass of all other objects change. It has always been clear that we would like to do better; the kilogram was the last SI unit still defined by a physical object. A reference defined with respect to a universal constant is not just more consistent, but also more accessible and practical. An idiosyncratic reference is, on the other hand, not very precisely shareable. Few people have access to the reference mass (there are six copies [2, 3]) and it is challenging to replicate (each has uniquely lost and gained atoms). While a universal reference is the ideal, there are tradeoffs between utility, universality and practicality that must be considered.

### The burden of success

What would an “ideal” reference genome look like? Because standards can take many forms, picking one is non-trivial. In practice, references can be a single sample or type, an average form or an empirical sampling, or a (universal) gold-standard. A major intent behind the original sequencing of the human genome was to provide a reference for future analysis and this has been wildly successful. The current reference genome works primarily as the backbone for all genomic data and databases. It provides a scaffold for genome assembly, variant calling, *-seq alignment, gene annotation and functional analysis. Genes are referred to by their loci, with their base positions defined by reference genome coordinates. Variants and alleles are labelled as such when compared to the reference (i.e., reference vs alternative). Diploid and personal genomes are assembled using the reference as a scaffold, and RNA-seq reads are typically mapped to the reference genome.

These successes make the reference genome an essential resource in many research efforts. However, a few problems have arisen:

1) The reference genome is idiosyncratic. The data and assembly which resulted in the reference reflect a highly specific process operating on highly specific samples. There could be real anomalies in the reference and it would be hard to know.
2) The reference genome is frequently confused with a healthy baseline. Experts likely know better, but it is still quite common to hear even genetics researchers refer to “errors in the reference” when they simply mean that the reference contains a minor allele. The tendency for this confusion to occur reflects a failed expectation for what a useful reference genome should look like.
3) The reference genome is hard to re-evaluate. Using a reference of any type imposes some costs and some benefits. Different choices will be useful in different circumstances but these are very hard to establish when the choice of reference is largely arbitrary. If we pick a reference in a principled way, then those principles can also tell us when we should not pick the reference for our analyses.

In the following, we briefly address these three points by outlining the history of the human reference genome, demonstrating some of its important properties, and describing its utility in a variety of research ecosystems. Finally, we suggest a route toward its improvement through the use of a consensus genome.

## The reference genome is idiosyncratic

### The history of the human reference genome

It is commonly said that we now live in the age of ‘Big Data’. In genomics, this refers to the hundreds of thousands of a genomes sequenced from across all domains of life (among grander plans [4]). The number of base pairs deposited in databases dedicated to sequencing data alone is at peta scale (e.g., Short Read Archive stands at ~2 x 10^16^ bp). This all started humbly enough with advent of Sanger sequencing in 1977. With the ability to read out the genome at base pair resolution, researchers were able to access the genetic code of bacteriophages and their favorite genes. Why sequence the full human genome, or any genome for that matter? The first reason was the desire for “Big Science” for biology [5]. Large projects existed in other fields such as physics, so why not biology? If other species were being sequenced, then why not humans? Of course there were more pragmatic reasons for the suggestion. In addition to demonstrating technological feasibility, genome-scale science would enable comprehensive investigation of genetic differences both within and across species [6, 7]. Additionally, sequencing an entire genome would allow for the identification of all genes in a given species, and not only those that were the target of a monogenic disease (e.g., *HTT* in Huntington’s disease [8]) or of interest to a field (e.g., *P53* in cancer [9]). The sequences of genomes would serve as useful toolboxes for probing unknown genomic regions, allowing for functional annotation of genes, the discovery of regulatory regions and potentially novel functional sequences. With these various desires in mind, the Human Genome Project was conceived [10].

The Human Genome Project was a gargantuan effort for its time, costing close to 3 billion US dollars on its completion. The first draft genome was published in 2001 [11], along with the competitive project from Celera [12]. The “complete” genome was announced in 2003, meaning 99% of the euchromatic sequence, with multiple gaps in the assembly [13]. Along with launching the field of human genomics, the project also prompted the development of many of the principles behind public genomic data sharing through the Bermuda Principles, ensuring the reference genome was a public resource [14]. As a direct consequence, the use and improvement of the reference has made genomics a rapidly growing and evolving field. The first major discovery was the scale at which the human genome was littered with repetitive elements, making both sequencing hard and the assembly of the sequenced reads a computationally challenging problem [15]. In time, single-molecule technologies generating longer reads [16–18] and algorithmic advancements [19–21] have been able to significantly improve the reference. Currently, the human genome is at version 38 (GRCh38 [22]), with less than 1000 reported gaps [23, 24].

## The reference genome is not a baseline

### The current reference genome is a type specimen

Although the reference genome is meant to be a standard, what that means in a practical sense is not clearly defined. For example, the allelic diversity within the reference genome is not an average of the global population (or any population), but rather contains long stretches highly specific to one individual. Of the 20 donors the reference was meant to sample from, 70% of the sequence was obtained from a single sample, ‘RPC-11’, who had a high risk for diabetes [25]. After the sequencing of the first personal genomes in 2007 [26, 27], the differences between genomes suggested the reference could not easily serve as a universal “background” genome. This observation is easily extended to other populations [28–31], where higher diversity can be observed. The HapMap project [32, 33] and the subsequent 1000 Genomes Project [34] were a partial consequence of the need to sample broader population variability [35]. Although the first major efforts to improve the reference focused on the need to fill in gaps (with other genomes), work is now shifting to incorporate diversity through the addition of alternate loci scaffolds and haplotype sequences [36]. But just how similar to a personal genome is the current reference? We performed a short series of analyses to test this (see **Figure 1**), using the 1000 Genomes Project samples. Starting off with allele frequencies (AF) of known variants, around 2 million reference alleles have population frequencies less than 0.5, indicating that they are the minor allele (dark blue line in **Figure 1A**). Of the total number of variants and base pairs assessed, this might seem high for a reference. In fact, the allelic distribution of the current reference is almost identical to the allelic distributions of personal genomes sampled from the 1000 Genomes Project (light blue lines in **Figure 1B**). The current reference is a well-defined (and well-assembled) personal genome. As such, it is a good type specimen, displaying properties of many personal genomes. However, this means it does not represent a default any more than any other arbitrarily chosen personal genome would.

**Figure 1.**
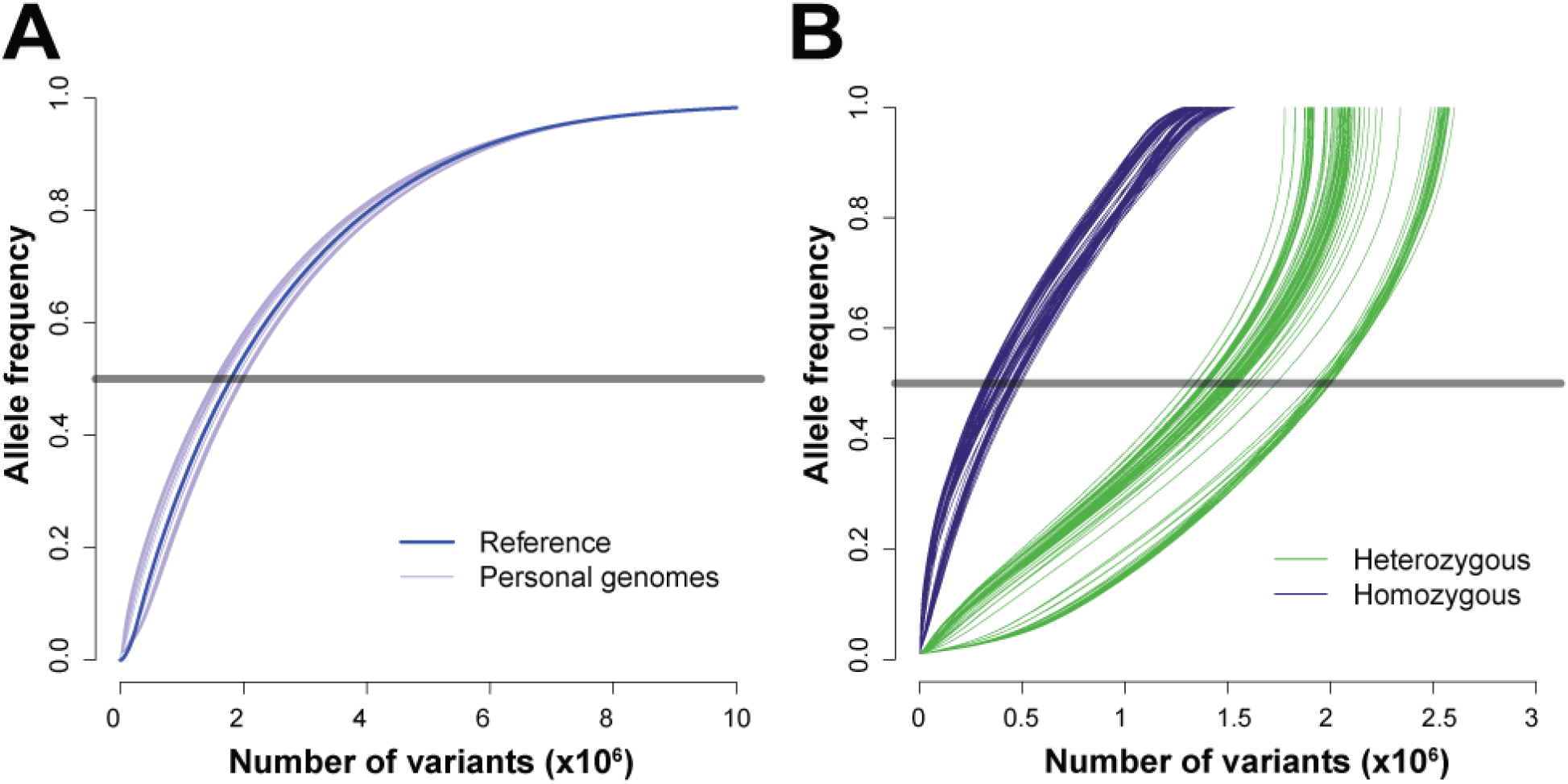
The reference genome is a type specimen. (A) Distribution of variants in the reference genome and those of personal/individual genomes. Collapsing diploid whole-genomes genotyped from the 1000 Genomes Project into haploid genomes, we can observe just how similar the reference is to an individual genome. First, taking population allele frequencies from a random sample of 100 individual genomes, we generated new haploid “reference” sequences. We replaced the alleles of the reference genome with the personal homozygous variant, and a randomly chosen one of the heterozygous alleles. For simplicity, all calculations are done against the autosomal chromosomes of the GRCh37 assembly and includes only single nucleotide bi-allelic (i.e., only 2 alleles per SNP) variants. (B) Distribution of allele frequencies for variants called in 100 randomly chosen personal genomes, computed against the reference genome. Here, variants with respect to the reference are quite likely to mean it is the reference which has the “variant” with respect to any default expectation, particularly if it is homozygous.

### Reference bias

Because the reference genome is close to being a type specimen, it can distort results where it is not very typical. In alignment, reference bias refers to the tendency for some reads or sequences to map more readily to the reference alleles, while reads with non-reference alleles may not be mapped or mapped at lower rates. For allele specific expression, eQTL analysis, and other RNA-seq based quantifications, the impact of this is uncertain [37–40]. In variant calling, this reference bias can be more important. Alignment to the reference to infer variation related to disease is still a step in most analyses, and crucial in clinical assignments of variant significance and interpretation [41, 42]. In these cases, reference bias will induce particular error. Variant callers can call more “variants” when the reference alleles are rare, or fail to call variants when they are rare yet also shared by the reference [43–46]. Due to the presence of rare alleles, some known pathogenic variants are easily ignored as benign [25]. A variant called with respect to the reference genome will be biased, reflecting properties of the reference genome rather than broadly shared population properties. Indeed, continuing with our analysis (**Figure 1B**), if we compare the variant calls within personal genomes against the reference, we find that close to ~⅔ of the homozygous variants (blue lines) and ⅓ of the heterozygous variants (green lines) actually have allele frequencies above 0.5. Variants with respect to the reference are quite likely to mean it is the reference which has the “variant” with respect to any default expectation, particularly if it is homozygous.

## The reference genome is hard to re-evaluate

### Type references are often good enough

A research ecosystem has grown up around the reference and has mostly taken advantage of its virtues while compensating for its flaws. In alignment for example, this has included the use of masked, enhanced, or diploid references. Masking repetitive regions or rare variants is a partial solution for improving mapping and assembly of short reads. Enhanced and diploid genomes involve inserting additional alleles or sequences into the current reference [45–53] which helps remove reference bias. And as the reference genome is a collapsed diploid, work on purely homozygous genomes (termed platinum references) will provide true haploid genomes (such as one from a molar pregnancy like the CHM1 cell line [54, 55]). More long term fixes are to generate new independent alternate references, eliminating the particularity of the original sample choices made such as those proposed by the MGI Reference Genome Improvement project [56]. The goal there is to amend the lack of diversity of the reference by creating gold genomes: gold-standard references each specific for an individual population. Along with latter being used as new standard genomes, personal or personalized genomes will become more common in clinical settings, with one’s own genome (potentially from birth), being used across one’s life for diagnostic assessments. However, the improvements all still use the reference genome as a foundation, in one form or another.

### Change is tricky

The pan-genome, a collection of multiple genomes from the same species, has been suggested to replace a single reference [57]. More complex than a single haploid reference sequence, it contains all possible DNA sequences which may be missing from any one individual [58]. A pan-genome can be represented linearly as concatenated alternate scaffolds, or as a directed graph [59], where alternate paths stand in for both structural and single variants [60]. These are particularly useful for plants where ploidy within a species exists [61], or in bacteria where different strains have lost or gained genes [62]. Adopting the graph genome as a reference reflects not just the inclusion of additional data, but is also a novel data structure and format. While well defined, it is non-trivial to incorporate into existing research practice and tools are under active development [63, 64]. A human pan-genome may improve variant calling by virtue of containing more variation [65], but this is offset by being a hard reference to refer to; unlike a linear reference genome, the coordinates in a pan-genome are harder to represent statically [66]. This is an issue as the current reference genome is the backbone of all genomics data. Variant databases use the reference’s coordinate systems, as do most gene and transcript annotations. Genome browsers use linear tracks of genomic data, and graph visualizations are unlikely to be interpretable (e.g., cactus graphs [67]). While graph genomes have many properties to recommend them, they will come at some cost and obtaining buy-in may be particularly challenging.

## Seeking consensus

### Why a consensus?

Instead of completely discarding the current reference or jumping to new graph genomes, we suggest an intuitive improvement – a consensus genome. In the same way that consensus sequences of transcription factor binding motifs represent the most common version of the motif, a consensus genome represents the most common alleles and variants of a population. A consensus genome is comparatively painless to existing research practice, looking substantially like a new reference in the current mode, but reflecting real improvements in interpretation and generalizability to new uses. In its very nature a consensus genome addresses the three concerns we have with the current reference: it is easy to replicate and is accessible, it is empirical and thus a baseline and is easily open to novel evaluation and adjustment to suit different baselines (e.g., populations).

### What would a consensus genome look like?

In the simplest of cases, a consensus genome remains a haploid linear reference, where each base pair represents the most commonly observed allele in a population. As a parallel to our assessment in the previous section, we show this by looking at the variants called for the personal genomes sampled from the 1000 Genomes Project (**Figure 2**). For illustrative purposes, we constructed a consensus genome by replacing all alleles with their major allele (**Figure 2A**), as measured in the 1000 Genomes Project dataset. Repeating the previous analysis, firstly we note the distribution of alleles are all above 0.5 as designed (**Figure 2B**). Secondly, the personal variants called are all below the population frequencies of 0. 5 as expected and we see that the total number of variants called has been significantly reduced (**Figure 2C**). Importantly, the number of homozygous variants called when using the consensus rather than the current reference is reduced from ~1.5M to ~0.5M. The distribution of the number of homozygous variants in all personal genomes in the 1000 Genomes Project collection against the standard reference (blue line) and consensus reference (red line) has markedly shifted (**Figure 2D**).

**Figure 2.**
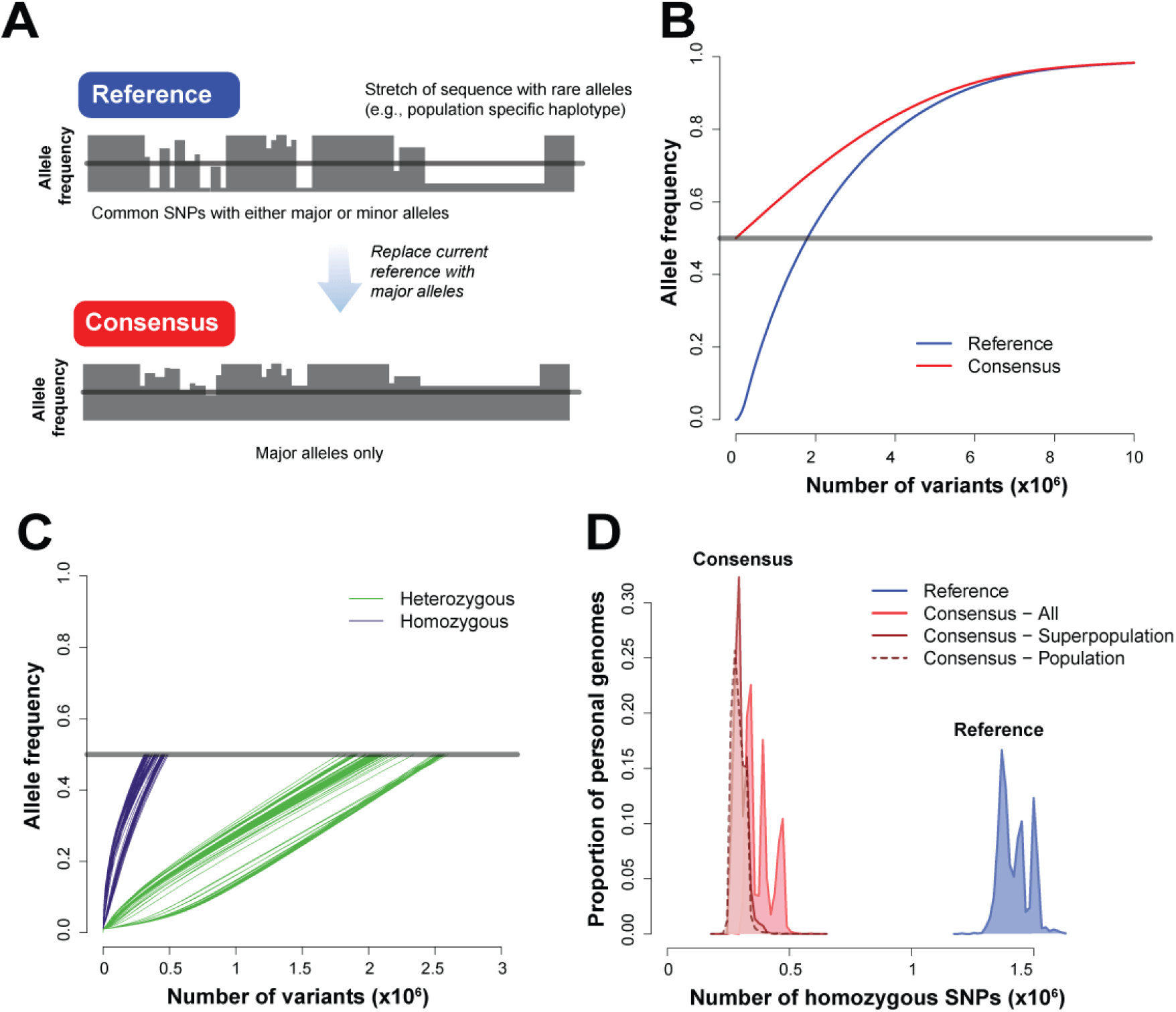
How consensus alleles improve the interpretability of the reference. (A) To build a consensus genome, we replace minor alleles within the current reference with their major alleles (AF>0.5) across all bi-allelic SNPs. (B) Distribution of variants in consensus (red line) and the current reference (blue line). (C) Distribution of allele frequencies for variants in 100 randomly chosen personal genomes, computed against a consensus genome. (D) Distribution of the number of homozygous SNVs in 2504 personal genomes, computed against the reference, against all-human consensus, mean of super-population consensuses and mean of population consensuses. The consensus reference for each of the 5 super-populations leads to additional reduction in the number of homozygous variants in the personal genomes in each super-population (dark red curve). Further breakdown into the 26 representative populations does not dramatically reduce the number of homozygous variants (dashed red line).

Additionally, the reference genome can be far from the average not just randomly (minor alleles) but also systematically, reflecting variation drawn from a particular population. A recent pan-assembly of African genomes directly spoke to the necessity for population specific references, as approximately 10% of DNA sequence (~300Mbp) was “missing” from the GRCh38 reference [68]. Indigenous and minor populations are understudied in general, which will need to be remedied in order to provide adequate clinical and medical care [69]. For example, certain drugs will be more effective and safer in some populations over others due to variants and how they change drug metabolism. To expand on this and test for population-specific impacts, we now build population-specific consensus genomes using allele frequencies of the five major populations represented in the 1000 Genomes Project data. Population-specific consensus genomes display a modest reduction in the number of homozygous variants called (darker red lines in **Figure 2D**), and a tightening of the spread of the distribution, as would be expected of a more refined null. This suggests the modal peaks are population specific variants, and use of population typical data is a boon.

### What would research built around a consensus genome look like?

In the above, our “consensus” uses both the existing reference and our knowledge of population allele frequencies. While this is particularly straightforward for SNPs, more complex genomic rearrangements can also be iteratively incorporated into a consensus genome. Practically speaking, any novel variant is called with respect to an existing reference and once that variant is known to be common, it becomes part of the new consensus. The distribution of population frequencies for variants means that relatively few genomes are necessary to ascertain that a novel variant is the majority allele. This makes the iterative improvement of the reference a more community-based effort, and one which can be tailored to suit different purposes. We think explicit choices of alternative references, particularly population-specific ones, will be a natural extension of the framework we describe (**Figure 3**). Switching to a consensus genome is not a transformational change to current practice and is far from a perfect standard, but by offering incremental, broad-based, and progressive improvement, we believe it is a timely step to take.

**Figure 3.**
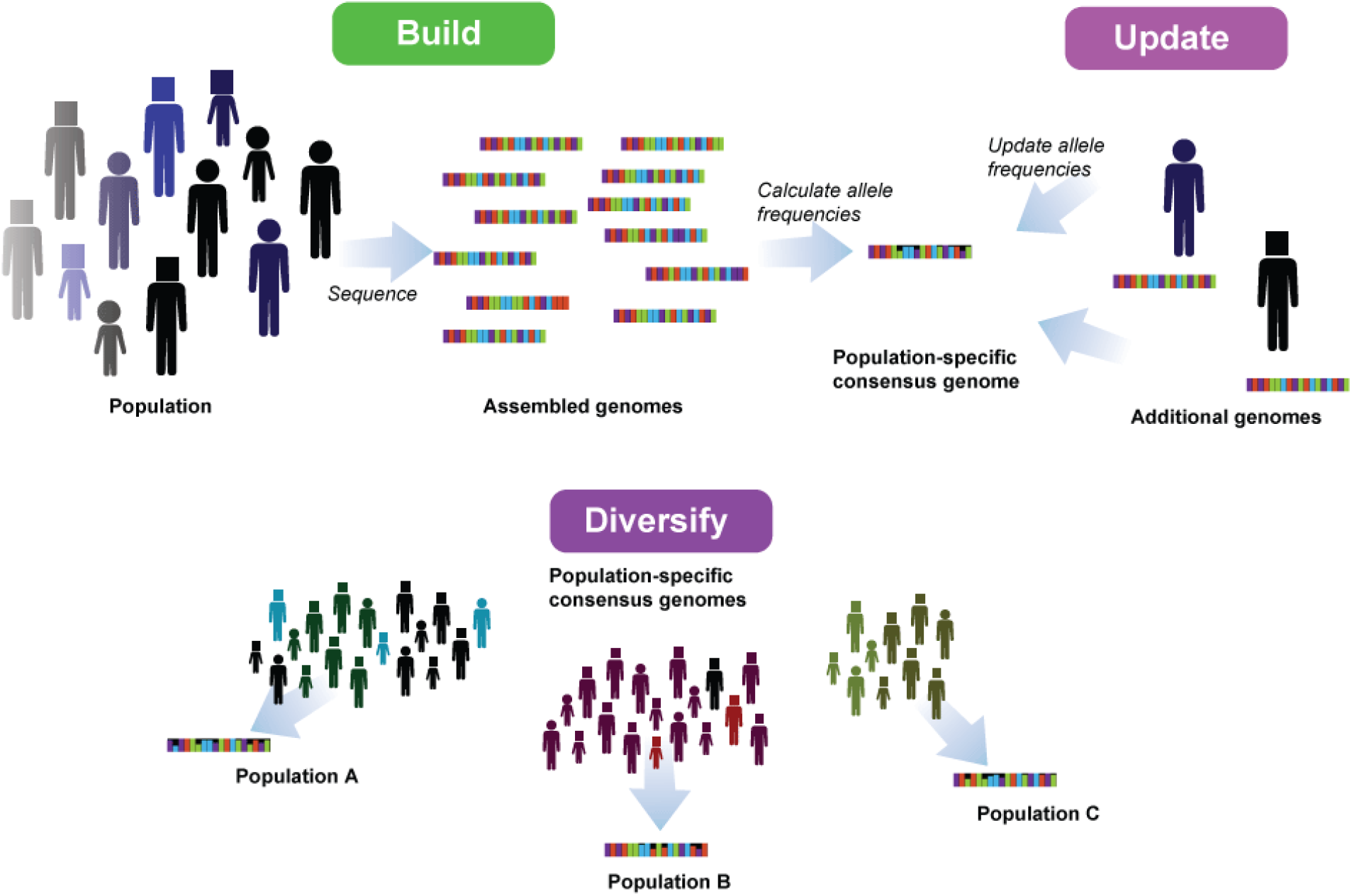
How-to reference. For future or new populations, sequencing is followed by building the consensus sequence from those genomes. Any new genomes will only adjust and improve on the current consensus based on change of allele frequencies. Finally, the reference can be replicated and diversified into other population-specific references.

## List of abbreviations

AF: allele frequency

## Declarations

### Ethics approval and consent to participate

Not applicable

### Consent for publication

Not applicable

### Authors’ contributions

AD ran the analyses. AD and JAG conceived the manuscript. SB, AD, and JAG wrote the manuscript. All authors read and approved the final manuscript.

## Acknowledgements

The authors would like to thank Megan Crow and Stephan Fischer for insightful comments and feedback on the manuscript.

